# Combining theory, model and experiment to understand how theta rhythms are generated in the hippocampus

**DOI:** 10.1101/115949

**Authors:** Katie A. Ferguson, Alexandra P. Chatzikalymniou, Frances K. Skinner

## Abstract

Scientists have observed theta rhythms (3–12 Hz) in the hippocampus for decades, but we do not have a clear understanding of how they are generated. This is largely due to the complex, multi-scale and nonlinear nature of the brain. To obtain insight into mechanisms underlying the generation of theta rhythms, we develop cellular-based network models of the hippocampus based on a whole hippocampus *in vitro* preparation that spontaneously generates theta rhythms. Building on theoretical and computational analyses, we find that spike frequency adaptation and post-inhibitory rebound constitute a basis for theta generation in large, minimally connected CA1 pyramidal (PYR) cell network models with fast-firing parvalbumin-positive (PV+) inhibitory cells. The particular theta frequency is more controlled by PYR to PV+ cell interactions rather than PV+ to PYR cell ones. We identify two scenarios by which theta rhythms can emerge and they can be differentiated by the ratio of excitatory to inhibitory currents to PV+ cells, but not to PYR cells. Only one of the scenarios is consistent with data from the whole hippocampus preparation, which leads to the prediction that the connection probability from PV+ to PYR cells needs to be larger than from PYR to PV+ cells. Our models can serve as a platform on which to build and develop an understanding of *in vivo* theta generation, and of microcircuit dynamics in the hippocampus.

**Significance:** Brain rhythms have been linked to cognition and are disrupted in disease. This makes it essential to understand mechanisms underlying their generation. Theory and mathematical models help provide an understanding and generate hypotheses. Together with experiment they contribute a framework to dissect the cellular contributions to network activity. However, models are inherently biological approximations, and thus the specific experimental and theoretical context upon which they are built will shape their output. If the approximations and contexts are not taken into account, particularly when using previously constructed models, misinterpretations can arise. Here, we use both theory and microcircuit models derived from a specific experimental context to provide insight into cellular-based mechanisms involved in theta rhythm generation in the hippocampus.

## Introduction

The goals of mathematical modeling in Neuroscience are many and varied. However, for any particular study, modeling goals must be clear as they guide our decisions during model development. Developed models can be used to generate new hypotheses and to investigate interactions across different scales. As we work to achieve an understanding of the brain, it is helpful to consider what is meant by an explanation in Neuroscience. In this regard, Aristotle’s doctrine of the four causes (*Falcon, 2015*)- the material cause, the formal cause, the efficient cause and the final cause, as described in *Brette* (*2013*) is useful. The efficient cause is what triggers the phenomenon to be explained, the material cause refers to the physical substrate of the phenomenon, the formal cause is the specific pattern responsible for the phenomenon (as would be represented by a mathematical model), and the final cause is the function of the phenomenon. Typically, as stated by Brette, theoretical approaches to Neuroscience tend to focus on formal and final causes, and experimental approaches to Neuroscience tend to focus on material and efficient causes. While all four causes may be needed to obtain a complete understanding, considering these four causes can serve to clarify modeling goals and how and why various mathematical models are developed and used. In turn, this can serve to enhance collaborative efforts in Neuroscience.

Electrical oscillations are hallmarks of the brain that are linked to normal and pathological functioning (*Buzsaki, 2006*). Thus, it is essential to understand the mechanisms underlying their generation. A large part of the challenge in obtaining mechanisms underlying oscillation generation is due to the multi-scale nature of our brains with its biological complexity and cellular specifics (*Cohen and Gulbinaite, 2014*). Various ‘building blocks’ such as post-inhibitory rebound have long been known, as identified from individual neurons and small circuits (*Gjorgjieva et al., 2016*). However, it is unclear what the building blocks for oscillation generation in the mammalian brain might be. A dominant oscillation in the hippocampus is the theta (3–12 Hz) rhythm (*Buzsaki, 2002; Colgin, 2013, 2016*). These rhythms are associated with memory processing and spatial navigation, present when the animal is actively exploring or during REM sleep. In the human hippocampus, theta rhythms are linked to similar behaviours (*Lega et al., 2012*). It may also be the case that theta rhythms in humans are associated with a wider behavioural repertoire relative to rodents, as they are present without sensory input (*Qasim and Jacobs, 2016*). Although Jung and Kornmuller discovered theta rhythms almost eighty years ago in the rabbit (*Colgin, 2013*), how theta rhythms are generated is not yet clear.

To understand theta generation it is important to distinguish between specific brain structures and excitatory and inhibitory cell interactions within these structures. As discussed by Colgin (2013), it is traditionally thought that the medial septum (MS) is critical for the generation of theta since theta rhythms are disrupted when the MS is lesioned or inactivated. However, the hippocampus can exhibit theta rhythms without the MS (*Goutagny et al., 2009*). In addition, distinct inhibitory cell populations, such as parvalbumin-positive interneurons, fire at unique phases of the theta rhythm, and likely play an important role in their generation (*Varga et al., 2014; Amilhon et al., 2015*) To understand the varied functional roles of these dominant rhythms and how they are modulated and controlled, we need to clearly decipher the cellular mechanisms underlying their generation. There is always some balance of interacting constitutive elements. From a mathematical modeling perspective, this reduces to deciding what ‘parameters, parameters, parameters’ (*Skinner, 2012*) and values to use and whether and how to represent the biological system given that any mathematical model is an approximation of the biology.

In this paper, our goal is to develop microcircuit models that we can use to understand how theta rhythms are generated. We take advantage of theoretical insights, an *in vitro* whole hippocampus preparation that spontaneously expresses theta rhythms, and the ability to readily do thousands of network simulations with our developed mathematical models. We present an explanation for theta generation that has elements of efficient, material and formal causes, and suggest that essential building blocks are spike frequency adaptation and post-inhibitory rebound.

## Materials and Methods

Here we summarize our overall strategy and describe the experimental context of the whole hippocampus preparation and our developed mathematical models and analyses. We also describe previous and motivating modeling work that the results are built upon.

### Overall strategy

Our goal is to develop experimentally motivated microcircuit models of a hippocampal CA1 network to provide insight into the mechanisms underlying theta rhythm generation. Our approach is shown in the schematic of **Figure 1** where orange and black arrows refer to links in the present or previous work respectively.

**Figure 1.**
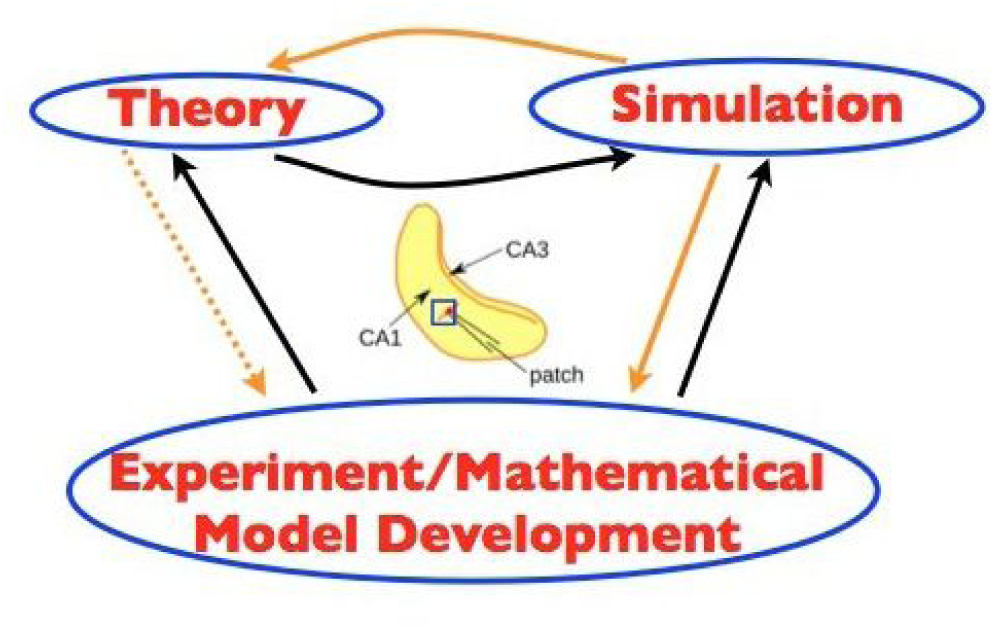
Overall Strategy. The three schematic parts (left, right, lower) of *Theory, Simulation* and *Experiment/Mathematical Model Development* are bidirectionally linked by arrows. *Theory* refers to mean field theory that was used to constrain the parameter sets to examine in simulations, using cellular models derived from experiment. *Simulation* refers to the computation of thousands of network simulations done. *Experiment/Mathematical Model Development* refers to the cellular, Izhikevich-type models that were developed in the experimental context of the whole hippocampus preparation. In the middle, a schematic of the whole hippocampus preparation with a an added blue square to illustrate the piece of tissue from the CA1 region of the hippocampus that is being modelled. The hippocampus schematic is adapted from Fig.1 of *Huh et al* (*2016*). Orange arrows are for links in the present work and black arrows for previous work. Dashed orange arrow from *Theory* to *Experiment* is because it is indirect as *Theory* previously contributed to *Simulation.*

We previously developed cellular, mathematical models of excitatory and inhibitory cells based on whole cell patch clamp recordings from the whole hippocampus preparation (*Ferguson et al., 2015b, 2013*). We used these cellular models to generate either excitatory (*Ferguson et al., 2015a*) or inhibitory (*Ferguson et al., 2013*) network models with sizes and connectivities as appropriate for the experimental context, and took advantage of a mean field theory (MFT) approach to determine parameter regimes in the excitatory networks (*Ferguson et al., 2015a*).

In the present work, we combine these excitatory and inhibitory networks and perform a detailed computational analysis of this network. We investigate the dynamic interplay between these two cell populations and their roles in theta generation. Further theoretical analyses are required to fully understand the network dynamics.

Overall, our strategy combines experiment, model development, simulation and theory. Our models bring together network size, connectivity and cellular characteristics in a closed fashion given the experimental context.

### Experimental context

In 2009, Goutagny and colleagues (*Goutagny et al., 2009*) developed an *in vitro* whole hippocampus rodent preparation that spontaneously generates theta (3–12 Hz) rhythms in the CA1 region. By blocking transmission across the septo-temporal axis, they identified multiple theta oscillators in the hippocampus (see Supplementary Fig.11 in *Goutagny et al.* (*2009*)). Further, the presence of these theta oscillations was not dependent on the CA3 region (see Supplementary Fig.10 in *Goutagny et al.* (*2009*)), and required *GABA_A_* and *AMPA* receptors (see Supplementary Table 1 in *Goutagny et al.* (*2009*)). Given this, we estimate that the minimum circuitry required for CA1 theta rhythms is contained in approximately 1 *mm*^3^. Using known cell densities and approximate volumes of axonal innervation (*Aika et al., 1994; Sik et al., 1995; Jinno and Kosaka, 2006; West et al., 1991; Hosseini-Sharifabad and Nyengaard, 2007*), we approximate that 30,000 excitatory, pyramidal (PYR) cells and 500 parvalbumin-positive (PV+) cells are involved in the spontaneous generation of theta rhythms in the CA1 region of the hippocampus. This size estimate is thoroughly described in our previous modeling work (*Ferguson et al.*, *2013*, *2015a*).

**Table 1.** Cell Model Parameters.

| Parameter | PYR |  | PV+ |
| --- | --- | --- | --- |
|  | weakly adapting | strongly adapting |  |
| $v_r$ (mV) | -61.8 | -61.8 | -60.6 |
| $v_t$ (mV) | -57.0 | -57.0 | -43.1 |
| $v_{peak}$ (mV) | 22.6 | 22.6 | -2.5 |
| $a$ (ms<sup>-1</sup>) | 0.00008 | 0.0012 | 0.1 |
| $b$ (nS) | 3 | 3 | -0.1 |
| $c$ (mV) | -65.8 | -65.8 | -67 |
| $d$ (pA) | 5 | 10 | 0.1 |
| $k_{low}$ (nS/mV) | 0.5 | 0.1 | 1.7 |
| $k_{high}$ (nS/mV) | 3.3 | 3.3 | 14 |
| $C_m$ (pF) | 300 | 115 | 90 |
| $I_{shift}$ (pA) | -45 | 0 | 0 |

Subsequent work by *Amilhon et al. (2015*) indicates that networks of PV+ and PYR cells could encompass the basic (minimal) units required for theta rhythm generation in the whole hippocampus preparation. This is because optogenetically silencing PV+ cells eliminates the theta rhythm, whereas silencing somatostatin-positive inhibitory cells do not. Further, it is the case that PV+ cells receive large excitatory postsynaptic currents (EPSCs) relative to PYR cells during ongoing theta rhythms. Also, simultaneous recordings from PV+ or PYR cells with the extracellular held recording of the theta rhythm indicates that the majority of PV+ cells are phasically firing with the rhythm, whereas PYR cells fire sparsely (*Huh et al., 2016*). We estimate that 20% or less of the PYR cells are firing based on (*Huh et al., 2016*)). Given that PV+ cells receive large EPSCs and that PYR cells fire sparsely, ongoing theta rhythms must necessarily be dependent on a large network effect.

During the ongoing theta rhythm, EPSCs in PYR cells are very small and variable (estimated to be less than 20 pA), whereas the inhibitory postsynaptic currents (IPSCs) to the PYR cells are larger (estimated to be approximately 200 pA). Conversely, the PV+ interneurons receive very large EPSCs (estimated to be up to 1000 pA), and smaller IPSC (approximately 200 pA). These estimates are based on whole cell current recordings from *Huh et al. (2016*). Given these estimates, EPSC/IPSC ratios for PYR cells are less than 1, and EPSC/IPSC ratios for PV+ cells are greater than 1.

Overall, we aim to determine the conditions under which our network models can produce population bursts at theta (3–12 Hz) frequency, given that there is sparse firing of excitatory PYR cells and non-sparse firing of inhibitory PV+ cells, and to capture the excitatory/inhibitory balance that is seen experimentally (*Huh et al., 2016*).

### Mathematical models

#### Cell Model

Our previously developed cellular models are based on experimental data from the *in vitro* whole hippocampus preparation (*Ferguson et al.*, *2013*, *2015b*). They are based on the mathematical model structure developed by Izhikevich (*Izhikevich*, *2010*, *2006*), in which the subthreshold behaviour and the upstroke of the action potential is captured, and a reset mechanism to represent the spike’s fast downstroke is used. Despite being relatively simple, parameter choices can be made such that they have a well-defined (albeit limited) relationship to the electrophysiological recordings. It has a fast variable representing the membrane potential, *V* (*mV*), and a variable for the slow “recovery” current, *u* (*pA*). We used a slight modification to be able to reproduce the spike width. The model is given by:

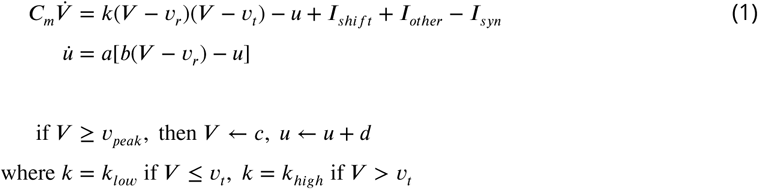

where *C_m_* (*pF*) is the membrane capacitance, *v_r_* (*mV*) is the resting membrane potential, *v_t_* (*mV*) is the instantaneous threshold potential, *v_peak_* (*mV*) is the spike cut-off value, *I_shift_* (*pA*) is a current that shifts the f-I curve laterally to allow the model to easily capture the rheobase current (for the strongly/weakly adapting models, rheobase current is 0/5 pA respectively), *I_syn_* (*pA*) represents the synaptic input from the presynaptic cell population (further details below), *I_other_* (*pA*) is an (excitatory) current drive to the network that is not directly modeled through *I_syn_* (further details below), *a* (*ms^−^*^1^) is the recovery time constant of the adaptation current, *b* (*nS*) describes the sensitivity of the adaptation current to subthreshold fluctuations - greater values couple *V* and *u* more strongly resulting in possible subthreshold oscillations and low-threshold spiking dynamics, *c* (*mV*) is the voltage reset value, *d* (*pA*) is the total amount of outward minus inward currents activated during the spike and affecting the after-spike behaviour, and *k* (*nS/mV*) represents a scaling factor. Parameter values for the cell models - strongly and weakly adapting PYR, and PV+ cell models are given in **Table 1**.

#### Synaptic Model

Synaptic input is modelled through a chemical synapse represented by:

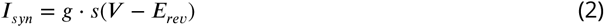

where *g* (*nS*) is the maximal synaptic conductance of the synapse from a presynaptic neuron to the postsynaptic neuron, *E_rev_* (*mV*) is the reversal potential of the synapse, and *V* (*mV*) is the membrane potential of the postsynaptic cell. The gating variable, *s*, represents the fraction of open synaptic channels, and is given by first order kinetics (*Destexhe et al. (1994)*, and see p.159 in *Ermentrout and Terman (2010)*):

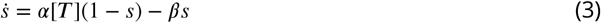

The parameters *α* (in *mM*^−1^*ms*^−1^) and *β* (in *ms*^−1^) in Equation 3 are related to the inverse of the rise and decay time constants (*τ_R_, τ_D_* in ms). [*T*] represents the concentration of transmitter released by a presynaptic spike. Suppose that the time of a spike is *t* = *t*_0_ and [*T*] is given by a square pulse of height 1 *mM* lasting for 1 *ms* (until *t*_1_). Then, we can represent

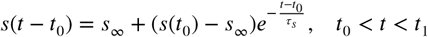

where

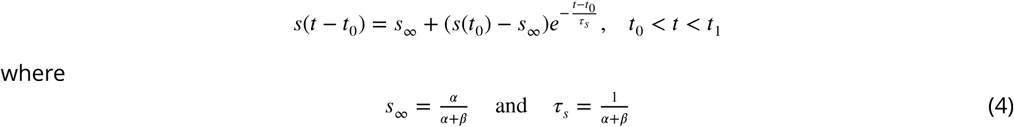

After the pulse of transmitter has gone, *s*(*t*) decays as

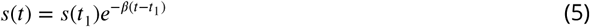

#### Network Models

Excitatory, inhibitory and excitatory-inhibitory network models are illustrated in **Figure 2** for PYR cell networks (top), PV+ cell networks (middle) and PYR-PV+ cell networks (bottom). Networks have either deterministic or noisy ‘other input’. If deterministic, then *I_other_* is a constant, tonic input to individual cells in the network where *I_other_* is chosen from a normal distribution with mean *I_app_*(*pA*) and standard deviation *σ_app_*(*pA*). If noisy, then *I_other_ = −g_e_*(*t*)(*V* −*E_rev_*). *g_e_*(*t*) is a stochastic process similar to the Ornstein-Uhlenbeck process as used by Destexhe and colleagues (*Destexhe et al., 2001)*

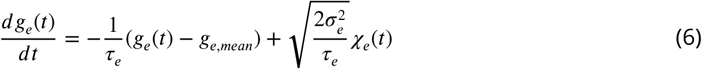

where *χ* _*e*_(*t*) is an independent Gaussian white noise processes of unit standard deviation and zero mean, *g_e,mean_* (*nS*) is the average conductance, *σ_e_* (*nS*) is the noise standard deviation value, and *τ_e_* is the time constant for excitatory synapses. *i*_*e*_ is fixed based on values as used in *Destexhe et al.* (*2001*) (*τ_e_ =* 2.73 *ms*).

**Figure 2.**
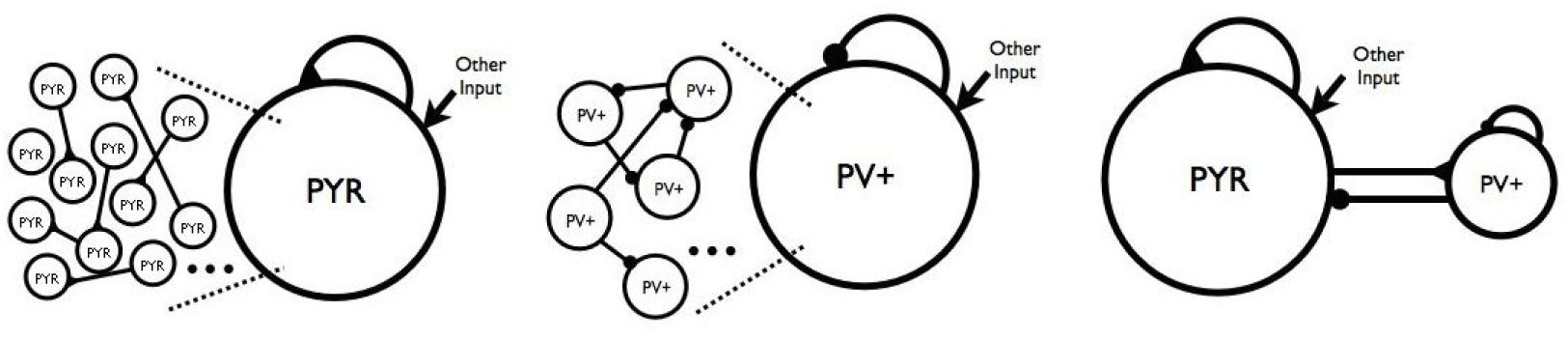
Network Schematics. (*left*) Excitatory, pyramidal cell networks. The large circle with PYR represents a population of individual pyramidal (PYR) cells and the black triangles represent excitatory connections. (*middle*) Inhibitory, fast-firing cell networks. The large circle with PV+ represents a population of individual inhibitory, parvalbumin-positive (PV+) fast-firing cells, and the black circles represent inhibitory connections. The connections are schematized so that there is clearly not all-to-all coupling. ‘Other input’ is either deterministic or noisy, excitatory synaptic input. (*right*) Excitatory-Inhibitory cell networks. PYR and PV+ cell populations combined to create networks of 10,500 cells. They are a combination of above schematics. However, now the excitatory input to PV+ cells comes directly from the PYR cell network.

#### Network Model Parameters, Rationale

Random connectivity was used throughout and the probability of connection is given in **Table 2** where it is fixed for PYR or PV+ cell networks, as estimated in previous work. Network sizes and synaptic time constants are given in **Table 2.** Excitatory and inhibitory reversal potentials *E_exc_*, *E_inh_* as derived from the experimental context are −15 and −85 mV respectively (*Huh et al., 2016*).

**Table 2.** Network Model Parameters. Connectivity between PV+ and PYR cells was not fixed, but ranged in values as indicated, A resolution of 0.01 was used from 0 to 0.1, and 0.1 upward.

|  | Number of cells | Probability of connection | Synaptic time constants |  |
| --- | --- | --- | --- | --- |
|  |  |  | Rise time (ms) | Decay time (ms) |
| **PV+** | 500 |  |  |  |
| PV-PV |  | 0.12 | 0.27 | 1.7 |
| PV-PYR |  | 0.01-1 | 0.3 | 3.5 |
| **PYR** | 10,000 |  |  |  |
| PYR-PV |  | 0.01-1 | 0.37 | 2.1 |
| PYR-PYR |  | 0.01 | 0.5 | 3 |

#### PYR cell networks

*g_pyr_* (nS) is the maximal excitatory (AMPA), synaptic conductance between PYR cells. Parameter explorations for deterministic networks included: *I_app_* (pA) = [0, 5,10,…, 75, 80], *σ_app_* (pA) = [0, 5,10, 15, 20]. Parameter explorations for noisy networks included: *g_e,mean_* (nS) = [0,1,2] (mainly), but values up to 10 explored, *σ_e_* (nS) = [0,0.2,0.4,0.6]. *g_pyr_* (nS) = [0.014, 0.024, 0.034,…, 0.084, 0.094] was used in both deterministic and noisy networks.

#### PYR-PV+ cell networks

A full exploration was done for connectivity between PV+ and PYR cell networks (see Table 2). We use *c*_*PV,PYR*_ and *c*_*PYR,Pr*_ to refer to the probability of connection from PV+ to PYR cells or from PYR to PV+ cells respectively. *g*_*pv*_ is the maximal inhibitory, synaptic conductance between PV+ cells, and a parameter value of *3nS* was used based on our previous PV+ cell network modeling (*Ferguson et al., 2013*).

##### Deterministic networks

Full connectivity explorations were done for chosen parameter sets: [*g_pyr_,I_a_*_*pp*_*,σ_a_*_*pp*_] = [0.014, 0, 0], [0.014, 0,10], [0.024, 30, 0], [0.054, 5, 5], [0.054, 20, 20], [0.074, 75,15]. *g_pv_*_*–pyr*_(*nS*) is the maximal, inhibitory, synaptic conductance on PYR cells from PV+ cells. It was fixed at 8.7 nS, as approximated from IPSCs in *Bartos et al.* (2002), for most of the simulations. *G_pyr_–_pv_* (*nS*) is the maximal, excitatory, synaptic conductance on PV+ cells from PYR cells. A value of 1 nS was used for most of the simulations, as estimated from *Papp (2013*).

##### Noisy networks

Full connectivity explorations were done for: (i) *g_e,mean_* (nS) = [0,1, 2], *σ_e_* (nS) = [0, 0.2, 0.4, 0.6], with *g_pyr_* = 0.014 nS, and (ii) All *g_pyr_* values given above, *σ_e_* = [0, 0.2, 0.4, 0.6] with *g_e,mean_* = 0 nS. *g*_*pv*–*pyr*_=8.7 nS was used but additional simulations using values of [6, 6.5, 7 …., 11.5,12] were done for two chosen parameter sets with *g_e,mean_* = 0 nS; [*σ_e_, g_pyr_, c_PY R Pv_, c_PV, PYR_*] = [0.2, 0.084, 0.4, 0.5] and [0.6, 0.014, 0.02, 0.3]. Similarly, *g_pyr–pv_=*3 nS was used and scaled for network size (see below). Additional simulations were done for values of [0.5,1,1.5, 2,…, 5.5, 6] for the chosen parameter sets. Overall, close to 6,000 simulations were performed.

#### Analyses

For each network simulation, we define the population activity as the average membrane potential of all model cells. Then, using the fast Fourier transform (fft), the network frequency (*f _peak_* in Hz) is defined as the frequency at which there is a spectral peak in the overall population activity. In this analysis, we disregard the initial transient activity (500 *ms*).

We defined a population burst based on the distribution of spikes of the PYR cell network. To do so, the total number of spikes within a small bin width were summed, where the bin width was dependent on the average peak frequency: bin width = *int*[*p*_1 *_ *round*[(*p*_2 *_ *exp*(−*p*_3 *_ *f*_*peak*_ + *p*_4_) + *p*_5_)/*p*_1_]], where *p*_1_ = 2, *p*_2_ = 2.0264, *p*_3_ = 0.2656, *p*_4_ = 2.9288, *p*_5_ = 5.7907 such that for peak frequencies ranging from *f_peak_* = 3–12 *Hz*, the bin width ranges from ≈ 23 – 7 ms. In this way, the bin would be smaller for higher frequencies. Then, the total number of spikes per bin width was normalized to its maximum (excluding transient activity within the first 500 *ms*) so that networks with significantly different levels of activity could be compared. A moving threshold capturing approximately five cycles was set to be the mean plus 0.35 standard deviations of the local normalized distribution. Then, the burst was determined to be the midway point between the increase past threshold and the previous decrease past the threshold (with the requirement that these points are at least 1/(*f* _*peak* *_ 2.5) *ms* apart). If the difference between the peak to trough of the burst is less than 0.2, it is no longer considered to be a burst. A population burst is considered to be more robust if the power of the fft is larger or the normalized size of the PYR spike distribution is larger. We note that if the population burst is reasonably robust, then the burst frequency as determined from the fft is essentially the same as the inverse of the burst width.

We automated the categorization of our network output for the different parameter sets explored. Specifically, non-firing cases were considered when there were < 300 spikes per burst bin. If network burst frequencies were within theta frequency ranges, they were further examined to determine their stability. Bursts were considered to be stable if there were at least two occurrences of two consecutive amplitudes decreasing by more than 79%. For each burst, we determined the burst width, the number of cells that fired in the burst, and the total number of spikes in the burst. In this way, we can track these properties not only for the network as a whole, but determine how they change over time. This analysis was based on custom code written in MATLAB.

For each simulation we recorded EPSCs and IPSCs from PYR and PV+ cells and chose a subset to analyze. Specifically, we used peakfinder in MATLAB, and ignored any peaks that were below a value thresholded at an order magnitude less than the main peaks. The first second was not included in the calculations. We computed averages and standard deviations of 3 PV+ and 5 PYR cells and rounded them out to give the reported values.

Simulations were run using the Brian simulator (*Goodman and Brette, 2009*) on the GPC supercomputer at the SciNet High Performance Computing Consortium (*Loken et al., 2010*) The initial conditions of our membrane potentials (*V*) were chosen to be uniform random values from –55 to –65 *mV*. We used the forward Euler method for integration with a time step of 0.02 *ms*. PYR cell networks were simulated for 10 sec whereas PYR-PV networks were simulated for 4 sec. For noisy simulations, simulations were done with second order explicit Runge Kutta numerical integration, with a time step of 0.04ms. A subset of these simulations were also run with the forward Euler method and compared.

### Motivating modeling studies

#### Excitatory (PYR Cell) Networks, Deterministic

In previous work, we considered whether CA1 PYR cell networks on their own could generate theta rhythms (*Ferguson et al., 2015a*). Or more specifically, given CA1 PYR cell intrinsic properties, connectivity, and cell numbers, can one obtain theta frequency (3–12 Hz) population bursting as observed in the experimental context? This was directly addressed in (*Ferguson et al., 2015a*). Individual PYR models were based on whole cell recordings from the whole hippocampus preparation (*Ferguson et al., 2015b*). Cells exhibited either weak or strong adaptation (determined by how much their frequency changed over the course of a one second-long input), and post-inhibitory rebound (spiking after being released from a hyperpolarizing current), and our models captured these properties. We connected these PYR cell models in a network (see left schematic of **Figure 2** and model details above) and took advantage of mean field theory (MFT) to find parameter regimes in which the network exhibited theta frequency population bursts (*Ferguson et al., 2015a*). Due to a scaling relationship between cell number, connection probability and *g_pyr_* from the MFT, we were able to use 10,000, rather than 30,000 PYR cells in our network simulations.

Stable theta frequency population bursts emerged from these PYR cell network models without a phasic drive. Similar to what was predicted from the MFT simulations, larger mean excitatory drive, *I*_*app,*_ was required to obtain theta frequency bursts as the standard deviation of the drive, *σ_app_,* increased (*Ferguson et al., 2015a*). Burst frequencies were determined by three factors: the mean excitatory drive, *I_app_* (which increases the burst frequency as it increases), the recurrent synaptic strength, *g_pyr_* (which decreases burst frequency as it increases), and the standard deviation of the excitatory drive across the cells, *σ_app_* (which increased frequencies as it increased) (*Ferguson et al., 2015a; Ferguson, 2015*).

From the parameter sets explored, network output was automatically categorized (see specifics in above sections) such that non-firing, stable theta frequency bursts, unstable bursts, or other groupings were apparent. In Figure 3 we show three example outputs, one of which exhibits theta rhythms (bottom), another exhibiting unstable bursts (top) and the other asynchronous behaviour (middle). In the theta bursting parameter regimes, we analyzed our networks to determine how many PYR cells were firing (i.e., active) during the population bursts. We found that > 90% of PYR cells are active when theta rhythms are present. In **Table 3**, we show the minimal number of active PYR cells for each *σ_app_* value. Except for *σ_app_ =* 0, these minimal cases all have frequencies that are below theta, but still have about 50% of their 10,000 PYR cells being active. Also, the number of active PYR cells during bursts increased with increasing *I_app_* and also with increasing *g_pyr_*. These observations are from simulations that were performed using strongly adapting PYR cell models. Weakly adapting PYR cell models were also used, but specifics are not shown - we already know from our previous MFT study that more input is required to drive them to achieve theta frequency population bursts with the reduced cellular adaptation (*Ferguson et al., 2015a*).

**Figure 3.**
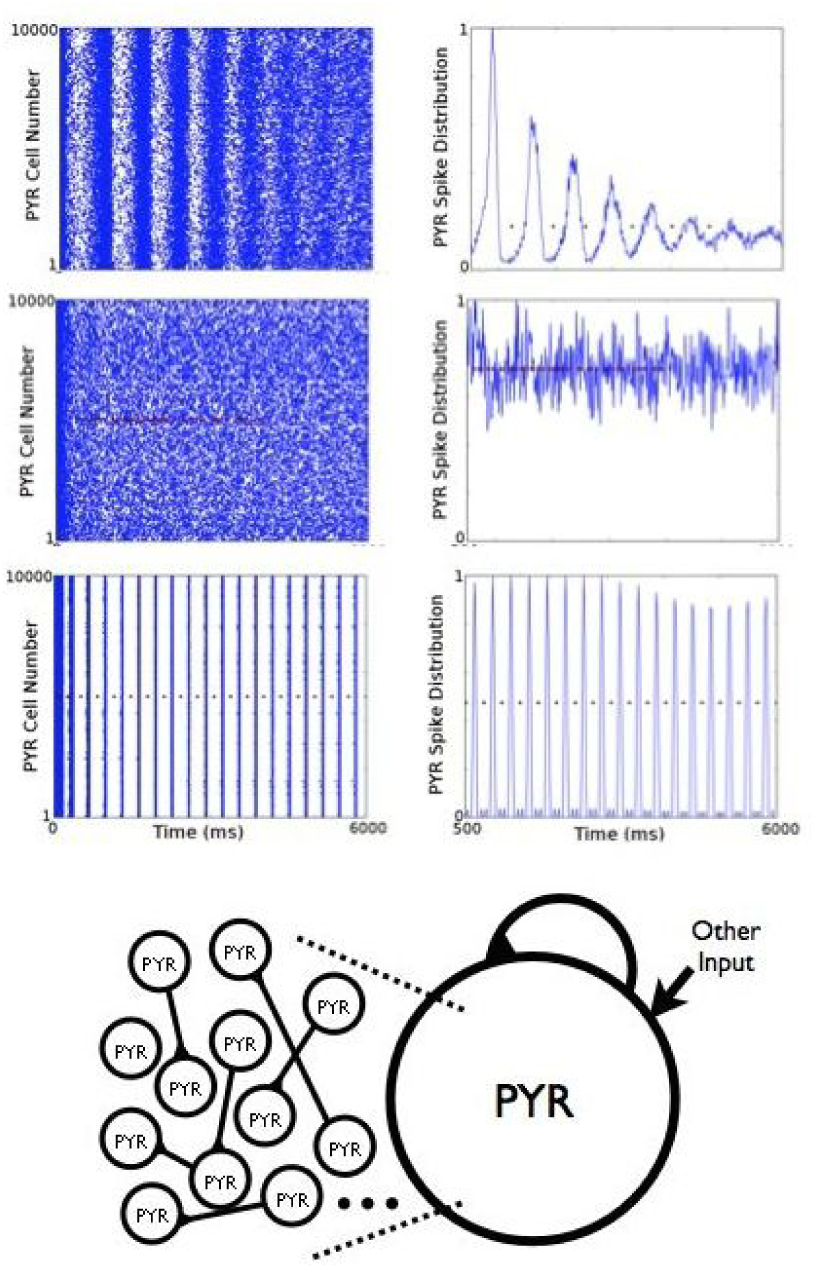
Deterministic Excitatory PYR Cell Networks. Three examples of raster plots and normalized spike distributions, one of which shows a theta frequency population burst. The spike distributions are used to determine the burst bins, and the red symbols represent the separation of the bins. *I*_*app*_ and ‘Other Input’ *Parameter values*: *I*_*app*_ = 0 *pA*, *g*_*pyr*_ = 0.014 *nS*, *σ*_*app*_ = 10 *pA* (*top - unstable bursts*); *I*_*app*_ = 0 *pA*, *g*_*pyr*_ = 0.014 *nS*, *o*_*app*_ = 10 *pA* (*middle* - *no bursts*); *I*_*app*_ = 30 *pA*, *g*_*pyr*_ = 0.024 *nS*, *σ* _*app*_ = 0 *pA* (*bottom* - 3.1 *HZ rhythm*).

**Table 3.** PYR Cell Networks: Numbers of Active, Firing Cells in Population Bursts.

| Parameters<br>$g_{pyr}(nS), I_{app}(pA), \sigma_{app}(pA)$ | Deterministic Networks | |
| --- | --- | --- |
|  | Average Number of Active Cells (/10,000)<br>per Population Burst | Burst Frequency (Hz)<br>(estimated from fft) |
| [0.064, 65, 0] | 9703 | 6.8 |
| [0.024, 10, 5] | 5541 | 1.4 |
| [0.054, 10, 10] | 4929 | 2.0 |
| [0.014, 10, 15] | 5075 | 1.2 |
| [0.014, 10, 20] | 5085 | 1.1 |
| $g_{pyr}(nS), I_{app}(pA), \sigma_{app}(pA)$ | Noisy Networks | |
|  | Average Number of Active Cells (/10,000)<br>per Population Burst | Burst Frequency (Hz)<br>(estimated from fft) |
| [0.014, 1, 0.6] | 5992 | 5.2 |
| [0.024, 1, 0.6] | 6173 | 5.2 |
| [0.034, 2, 0.6] | 6115 | 9.4 |
| [0.054, 2, 0.4] | 6757 | 9.4 |
| [0.074, 2, 0] | 6848 | 9.1 |

Thus, our models suggested that an appropriate balance between spike frequency adaptation and excitatory connectivity in CA1 PYR cell networks could provide an essential mechanism for theta population bursts. However, the majority of PYR cells in the models were active during the population bursts, which is *not* consistent with experiment. This led us to consider how inhibitory cells may contribute to burst dynamics.

#### Excitatory-Inhibitory (PYR-PV+ Cell) Networks, Deterministic

To build networks with both excitatory and inhibitory cells, we first take advantage of previous work in which we developed cellular models for PV+ fast-spiking cells based on experimental recordings from the whole hippocampus preparation (*Ferguson et al., 2013*). Given this work, estimates of EPSCs of approximately 1000 pA, and that we want our PV+ cells to be able to fire coherent bursts, we set *g_pv_* = 3 *nS* (*Ferguson et al. (2013*) and see also *Skinner and Ferguson (2013*)).

Now, rather than setting the excitatory drive to PV+ cell network as deterministic ‘other input’ as done previously in *Ferguson et al. (2013)* (see middle schematic of Figure 2), we create networks in which the excitatory drive comes directly from the 10,000 PYR cell network. This is shown in the right schematic of **Figure 2**. Since our model is designed to explore oscillatory activity intrinsic to the CA1 region of the hippocampus, input from other regions are not specifically included. We chose example PYR cell networks which exhibited distinct firing patterns (non-firing, stable bursts, etc.) and explored how the connectivity between PYR cells and PV+ cells affected network activity (*Ferguson, 2015*). This limited set of excitatory-inhibitory network simulations provided a motivating basis for the expanded set of simulations presented in the Results.

In PYR cell networks that exhibit stable bursts, introducing PV+ cells does not ensure the bursts are maintained, but instead requires that the connection probability from PV+ to PYR cells (*c_PV,P,YR_*) surpasses a critical value. This critical connectivity value depended on the PYR-PV+ connection probability (*c_PYR pV_*), as it could not be drastically lower than this value. Interestingly, if stable bursts are maintained, the frequency of the population bursts is always higher in the PYR-PV+ cell networks relative to the PYR cell networks alone. We note that for each set of simulations, we did not alter the excitatory drive to the PYR cells, and thus the cells did not increase their firing due to an external change in the amount of excitation. Rather, this increased burst frequency is due to the post-inhibitory rebound spiking of the PYR cells as a response to the inhibitory input from the PV+ cells.

Alternatively, if the original PYR cell networks were unstable or non-firing, stable population bursts in the theta frequency range could emerge with the inclusion of the PV+ cell population. In all cases, and in contrast with our PYR cell networks, it was possible to simultaneously obtain theta frequency population bursts and sparse firing of the PYR cells. These observations are illustrated in **Figure 4**. Thus, even when oscillations did not exist or were not stable in the PYR cell networks, the influence of PV+ cells could lead to stable network rhythm generation and sparse firing. However, this depended upon an appropriate balance of connection probabilities between the two populations.

**Figure 4.**
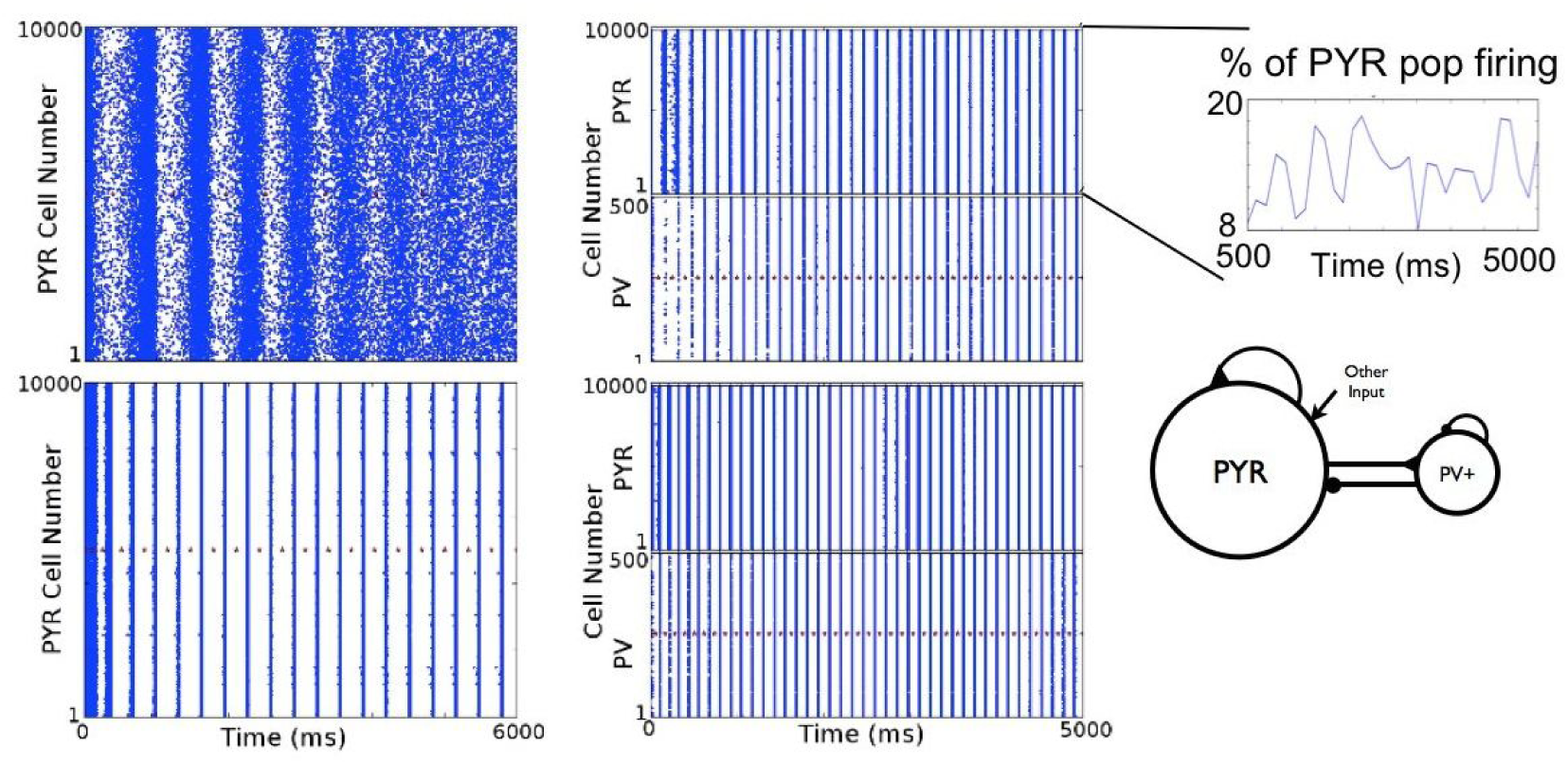
Deterministic Excitatory-inhibitory PYR-PV+ Cell Networks. Examples of population burst stabilization, sparse excitatory firing and network frequency increase are shown. PYR cell networks are shown on the *left* (same examples from *Figure 3*). PYR-PV+ cell networks are shown on the *right* with connectivity parameters *c_PY R,pV_,c_PV, pYR_* of (0.05,0.2) and (0.01, 0.2) for *top* and *bottom* respectively. Inset show that PYR cells firing is less than 20%. The population burst frequency increases from 3.1 to 6.7 Hz (*bottom*).

Overall, we found that PYR cell networks could exhibit coherent firing at various frequencies (≈ 0.5 – 6 *Hz*) but when theta frequencies were produced, essentially all PYR cells were recruited to fire in every cycle - a behaviour that is not consistent with what is seen in our experimental setting, where PYR cells sparsely fire. However, the inclusion of PV+ cells allowed sparse firing to emerge while maintaining theta bursts but of increased frequency, or generating theta bursts. Since we did not change excitation in any other way, a key contributor to this is post-inhibitory rebound firing. Thus, population bursts at theta frequency in the PYR-PV+ cell networks with sparse firing depend on a number of factors, including the amount of input that the cells are receiving at any given point in time. As such, the precise connection probabilities in the models is not the essence, but rather the relative balance between excitation and inhibition. More complete exploration and analyses are needed to untangle this.

## Results

We use network models that are minimal but at the same time are constrained by experiment. In this way, we reduce the uncertainty in choices for parameters and parameter values. Detailed reasoning and rationale for our choices are provided in the Methods. We examine whether it is possible to capture the experimental observations, and if so, what are the underlying mechanisms that allow this? To start to address this, we presented previous work and motivating modeling simulations using deterministic networks in the Methods. Although limited, these simulations showed that there is an intricate tangling of cellular properties and excitatory and inhibitory balances that underlie the generation of theta population bursts with sparse pyramidal (PYR) cell firing. That is, there are many interconnected factors. We now perform an expanded set of simulations using more biologically realistic input to obtain insight into possible underlying mechanisms.

### Noisy networks

#### Excitatory PYR Cell Networks

We perform simulations with noisy, excitatory input. That is, **‘**other Input**’** in **Figure 2** is given by a stochastic rather than a deterministic process (see Methods). Since we know that PYR cells do not receive large EPSCs (*Huh et al., 2016*), we can focus on small mean excitatory conductances That is, an EPSC of 20 pA into a PYR cell would give an excitatory conductance estimate of less than 1 nS, given excitatory reversal potential and resting voltage values. However, since some of the resultant EPSC would be due to recurrent connections, values greater than 1 nS should be considered, but do not need to be that much larger since PYR cell connectivity is minimal. As such, we consider *g_e,mean_* values of 0–2 nS to be a wide enough consideration (with *σ_e_* values < 1) to fully encompass the biological situation present in the whole hippocampus preparation.

We first note that, unlike the deterministic PYR cell network simulations, the patterns are more variable as would be expected. However, similar to the deterministic simulations, burst frequency increased with increasing excitatory drive, *g_e,mean_,* and decreased with increasing recurrent synaptic strength *g_pyr_*. There were no theta frequency bursts for *g_e,mean_ =* 0, but for all *g_e,mean_* = 1 or 2, there were theta frequency population bursts except for the largest *g*_*pyr*_ = 0.094 nS value when *g*_*e*,*mean*_ = 1, *σ*_*e*_ = 0.6 which was outside of the theta frequency range. Example output is shown in **Figure 5** when there are (bottom) or are not (top) population bursts. Also shown in **Figure 5** is the PYR cell spike distribution which was used to determine whether bursts were present (see Methods). Looking at the number of active PYR cells, it always exceeded 50% and **Table 3** shows specific numbers for the five lowest number of active PYR cells from the full set of simulations. For weakly adapting cells, no theta rhythms were obtained for *g_e,mean_* = 0 or 1 but could emerge for larger input values. Since this would be beyond EPSCs values observed in experiment, we did not do further detailed explorations using weakly adapting cells. Thus, similar to the deterministic PYR cell network simulations, theta population bursts are present but never with sparse firing of PYR cells in more realistic, noisy PYR cell networks.

**Figure 5.**
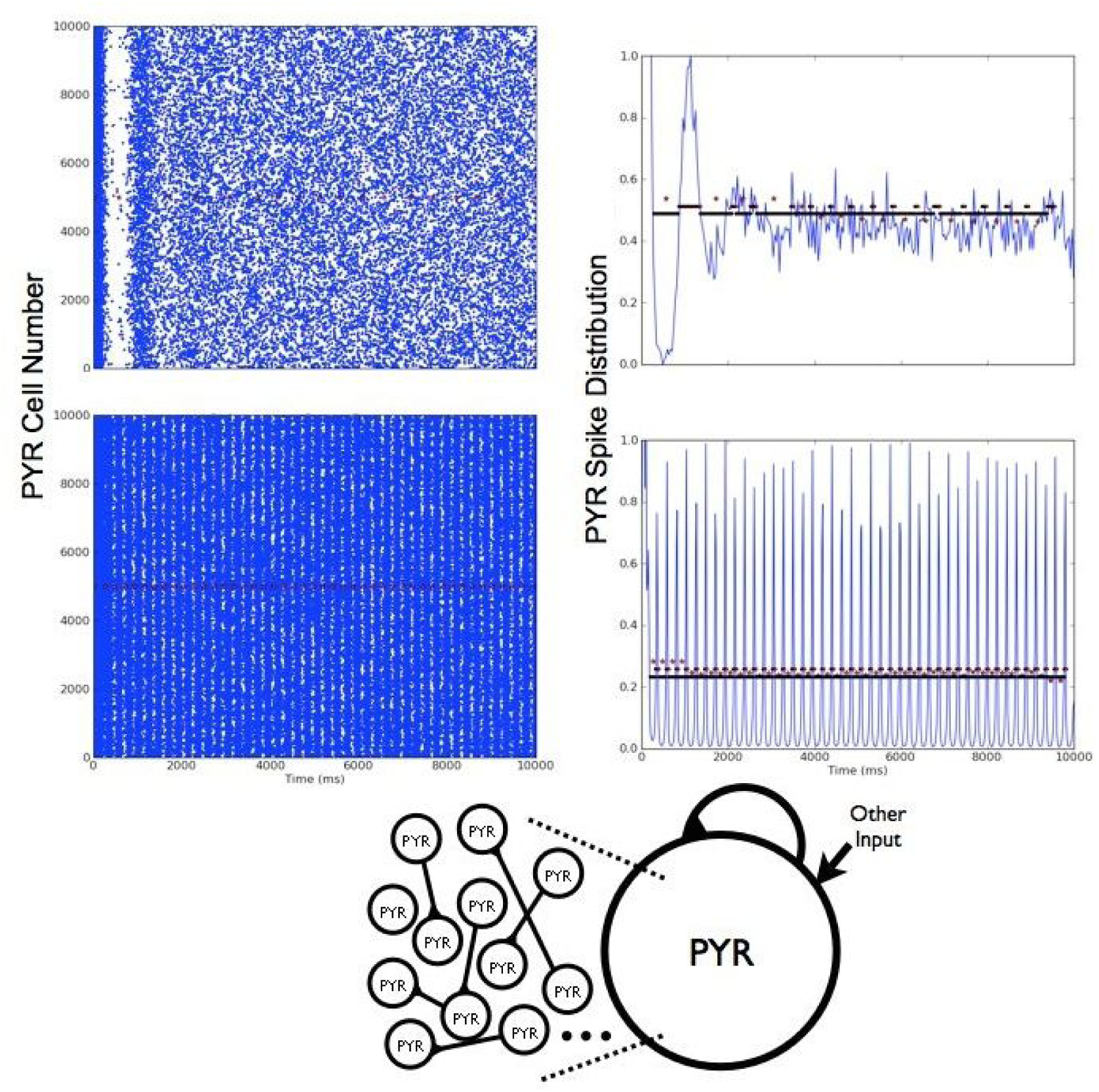
Noisy Excitatory PYR Cell Networks. Two examples in which theta rhythms (population bursts) are either present (*bottom*) or not (*top).* On the *left* is shown the raster plot, and on the *right* is shown the normalized spike distribution. Red stars delineate the automatic detection of separate bursts. *Parameter values: g_e,mean_ =* 0, *g_pyr_ =* 0.074, *σ_e_* =0.2 (*top row - no theta rhythm); g_e,mean_ =* 1, *g_pyr_ =* 0.074, *σ_e_ =* 0.2 (*bottom row - 4.4 HZ rhythm).*

#### Excitatory-Inhibitory PYR-PV+ Cell Networks

A full exploration of connectivities (*c_PV_, _PYR_* and *c_PY,R,PV_*) with *g_pyr_* = 0.014 and *g_e,mean_* = 0,1 and 2 nS was done. We first note that, similar to the deterministic simulations, theta bursts could be present in PYR-PV+ cell networks even if PYR cell networks did not have any population bursts. An example of this is shown in **Figure 6**. A summary of the burst frequencies are shown in the top part of **Figure 7** for *g_e,mean_* = 0 and 1 nS. Theta frequency bursts encompass colors that range from light blue to orange. Given this, it is clear that PYR-PV+ cell networks with *g_e,mean_* = 1 nS had burst frequencies that exceeded theta, except for when *σ_e_* = 0 nS, meaning noiseless networks. For *g_e,mean_* = 2 nS, the burst frequencies far exceeded theta frequencies (not shown). Also, since larger *g_pyr_* values result in an increased burst frequency (see **Figure 8**, right), it did not make sense to do additional simulations with *g_e,mean_* = 1 or 2 nS. This thus led to a focus on simulations with *g_e,mean_* = 0 nS. The full range of *g_pyr_* values were simulated along with explorations of PYR to PV+ and PV+ to PYR cell conductance values (see Methods).

**Figure 6.**
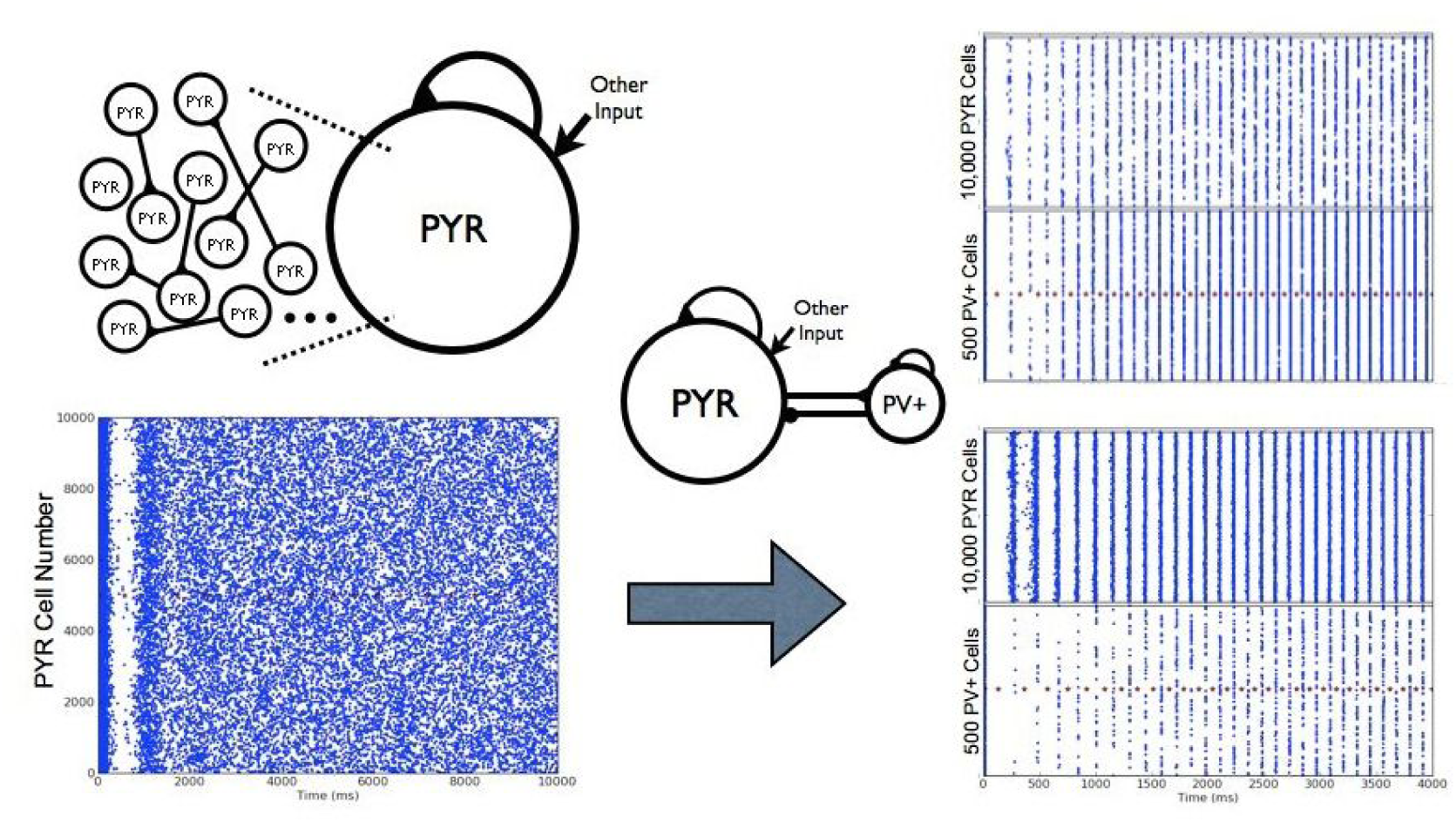
Noisy PYR to PYR-PV+ Cell Network Transition. From one of the PYR cell network examples in *Figure 5* where there are no theta bursts, two examples with different connectivities of PYR-PV+ cell networks are shown in which theta population burst rhythms emerge. Connectivity parameters *c_PY R,PV_,c_PV, PYR_* of (0.2, 0.5) and (0.02, 0.5) for *top* and *bottom,* respectively on the right. *Other parameter values: g_pyr_*(*nS*)*, g_e,mean_*(*nS*)*, σ_e_*(*nS*)*, g_pyr–pv_*(*nS*)*,g_pv–pyr_*(*nS*) = [0.074, 0, 0.2, 3, 8.7].

**Figure 7.**
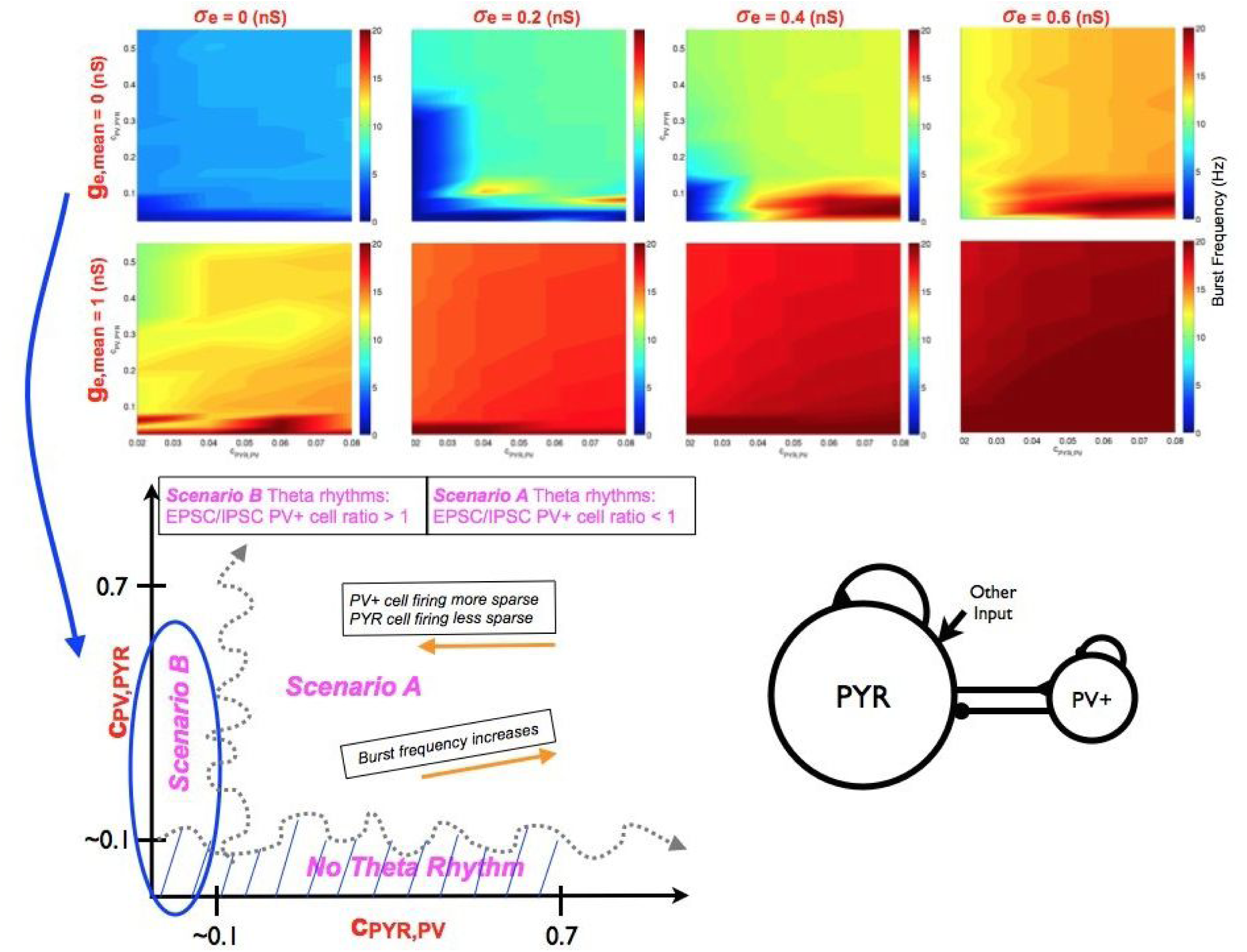
Parameter Balances for the Existence of Theta Rhythms in PYR-PV+ Cell Networks. (*Top*) Burst frequency summarized for a range of parameter values. Color plots shows how burst frequency changes for a range of input to the PYR cell population and for a range of connectivities between PYR and PV+ cell populations. Note that *c_PY,R,PV_* is shown for a much smaller range than ^*C*^*PV,PYR*. Moving horizontally, one sees that increasing *σ_e_* typically leads to an increase in burst frequency. It is also clear that beyond *g_e,mean_ =* 0, theta frequency rhythms are not present if *σ_e_* is non-zero. *Other parameter values: g_pyr_*(*nS*)*, g_pyr–pv_*(*nS*)*, g_pv–pyr_*(*nS*) =[0.014, 3, 8.7]. (*Bottom*) Theta rhythm generation overview. This schematic summarizes the balances in the generation of theta rhythms and their characteristics in the network models. Two scenarios, A and B, can be identified based on connectivity balances (*c_PV,PY R_* and *c_PY R,pV_*) and are approximately delineated by the squiggly grey dashed lines. Note that these separations are illustrative, as the exact connectivity boundary value will depend on the other parameters in the models. However, it is clear from the many simulations done and analyzed that one is able to differentiate these regions. Network models characteristics (frequency, and PV+ and PYR cell firings) are given in boxes with orange arrows. Scenario A and B are differentiated by their EPSC/IPSC ratios to PV+ cells, as given in magenta text.

**Figure 8.**
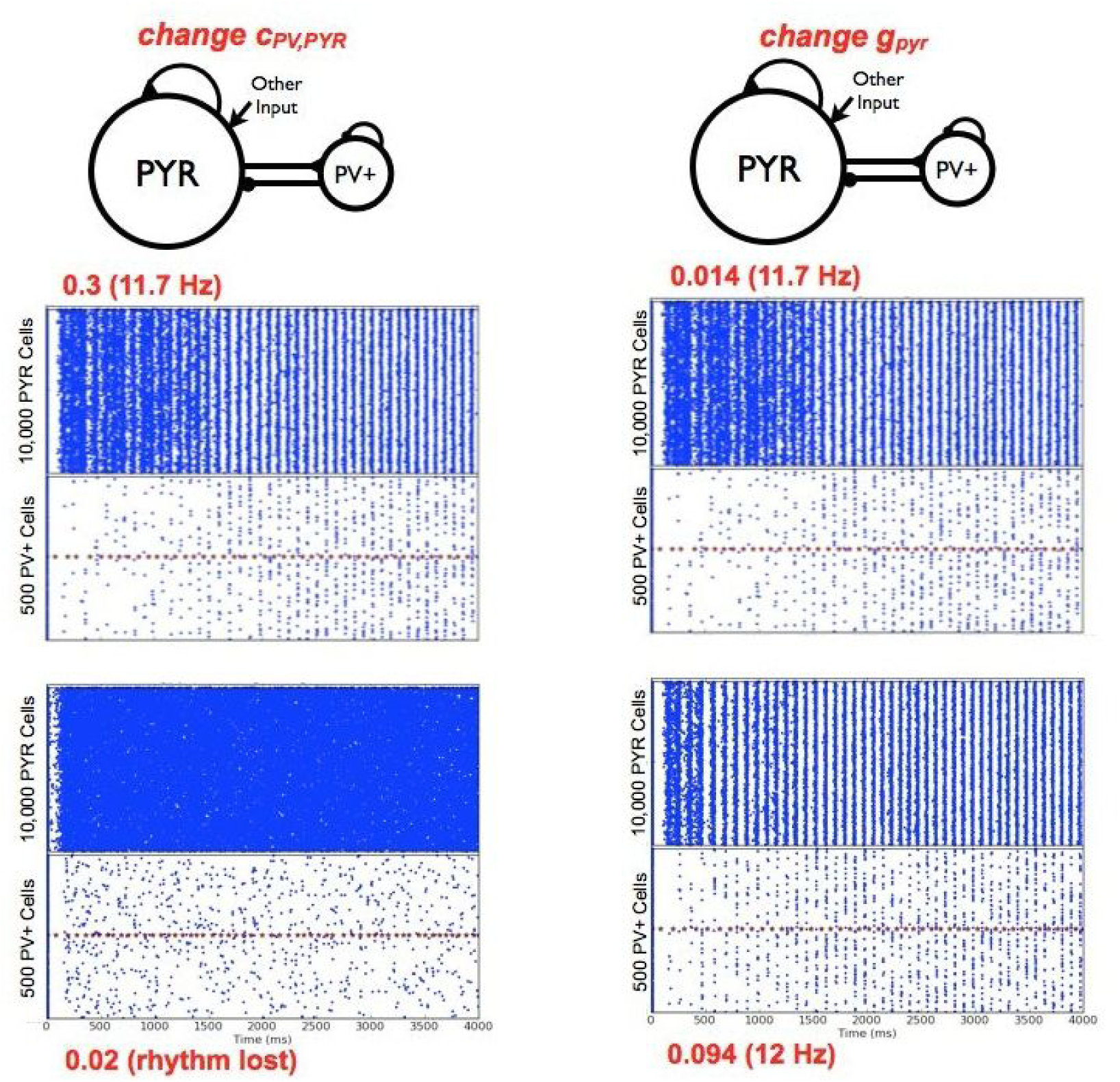
PYR-PV+ Cell Networks - Parameter Variations. (*Left*) Example raster plots for decreasing *C_PV,PYR_*. Two examples are shown, with the *C_PV, PYR_* value shown in red along with the network frequency as appropriate. The average number of cells firing per burst are 548 (PYR cells) and 22 (PV+ cells) for *C*_*PV*, *PYR*_ = 0.3. *Other parameter values: g*_*pyr*_ *g*_*e*,*mean*_, *σ*_*e*_, *g*_*pyr*−*pv*_, *g*_*pv*−*pyr*_, *c*_*PY R*,*PV*_ = [0.014, 0, 0.6, 3, 8.7, 0.02]. (*Right*) Example raster plots for changing *g_pyr_.* Two examples are shown, with the *g_pyr_* value shown in red along with the network frequency. The first, top example is the same as the one on its left. The average number of cells firing per burst are 514 (PYR cells) and 34 (PV+ cells) for *g_pyr_* = 0.094 nS. *Other parameter values: g_e_*,mean,*σ_e_*,*g_pyr–pv_*,*g_pv–pyr_*,*c_PY R,PV_*,*C_PV,PYR_ =* [0,0.6, 3, 8.7, 0.02, 0.3].

From a computational analysis that consisted of several thousands of simulations, we were able to obtain a ‘lay of the land’ in terms of required parameter balances for theta rhythms to occur, as well as their characteristics. This is schematized in the bottom of **Figure 7**. The connectivity ranges that refer to the summarized plots above are indicated. As can be seen, they are summarized only for smaller *c*_*PY R,PV*_ values. However, as *c*_*PY,R,PV*_ increases, the burst frequency increases further (not shown). As noticed in the motivating deterministic simulations, and similarly here for the noisy runs, if *c*_*PV,PYR*_ is too small, theta bursts are not present. That is, there needs to be enough connectivity from PV+ to PYR cells to have post-inhibitory rebound firing of an appropriate amount in PYR cells for theta bursts to occur. An example of how theta bursts are lost (when *c*_*PY,R,PV*_ is also small, is shown in **Figure 8**, left.

#### An essence of theta rhythm generation and experimental data ‘matching’

From our noisy and motivating deterministic simulations, we are able to distinguish two Scenarios, A and B, by which theta rhythms occur. It is due to the wide swaths of simulations and analyses of them that we are able to identify these different scenarios as shown in **Figure 7**. As schematized, the transition between scenarios is not meant to be literal, but illustrative to consider the several other parameters that would affect the exact transitions. The difference between these two scenarios lies in how active the PYR and PV+ cell populations are, which in turn affects the EPSCs and IPSCs; received by them.

So how do theta frequency bursts emerge in PYR-PV+ cell networks? We first point out that ‘ *g_e,mean_* = 0 nS with non-zero *σ_e_* is the appropriate ‘other input’ to use to have theta frequency population bursts. This is due to results from PYR cell networks and knowledge of EPSC values to PYR cells in experiment. Also, as noted above, if *c*_*PV,PY R*_ is not large enough, there will be no theta rhythms (**Figure 8**, left). This immediately indicates the importance of post-inhibitory rebound in PYR cells to generate theta bursts. However, while *c*_*PV, PY R*_ cannot be zero (since this would mean that PV+ cells are not coupled to PYR cells and one would have PYR cell network output, which we know has no theta bursts), the exact non-zero value needed for theta bursts to emerge also depends on *c*_*PY R,PV*_. The example shown in **Figure 8** (left) is for a small value of *c*_*PY R,PV*_ and the PV+ cells are not able to fire much or coherently. If *c*_*PV PYR*_ is decreased, but using a larger value of *c*_*PY R,PV*_, the PV+ cells are firing more, but the PYR-PV+ cell networks are not able to generate theta frequency bursts when *c*_*PV, PYR*_ is too small, presumably because now there is not enough post-inhibitory rebound to allow the PYR cells to exploit their spike frequency adaptation intrinsic properties and move it to a robust bursting in the theta frequency range. The importance of post-inhibitory rebound firing in the generation of theta rhythms was already suggested by *Goutagny et al. (2009)* from their experimental work, and so it is reassuring that the model results are consistent with this. Once one is within parameter balance regimes with robust theta frequency bursts, the frequency increases with increasing *c*_*PY, R,PV,*_ and less so for increasing *c*_*PV*, *PYR,*_ as determined from an overall examination of the simulations. This observation is particularly apparent in the upper summary plot of **Figure 7** (*g_e,mean_*=0 and *σ_e_*=0.6). This is illustrated by the slanted box (‘burst frequency increases’) in the schematic of **Figure 7**. The changing burst frequency can be seen if *g_pyr–pv_* rather than *c*_*PY R,PV*_ is modified, as shown in **Figure 9**.

**Figure 9.**
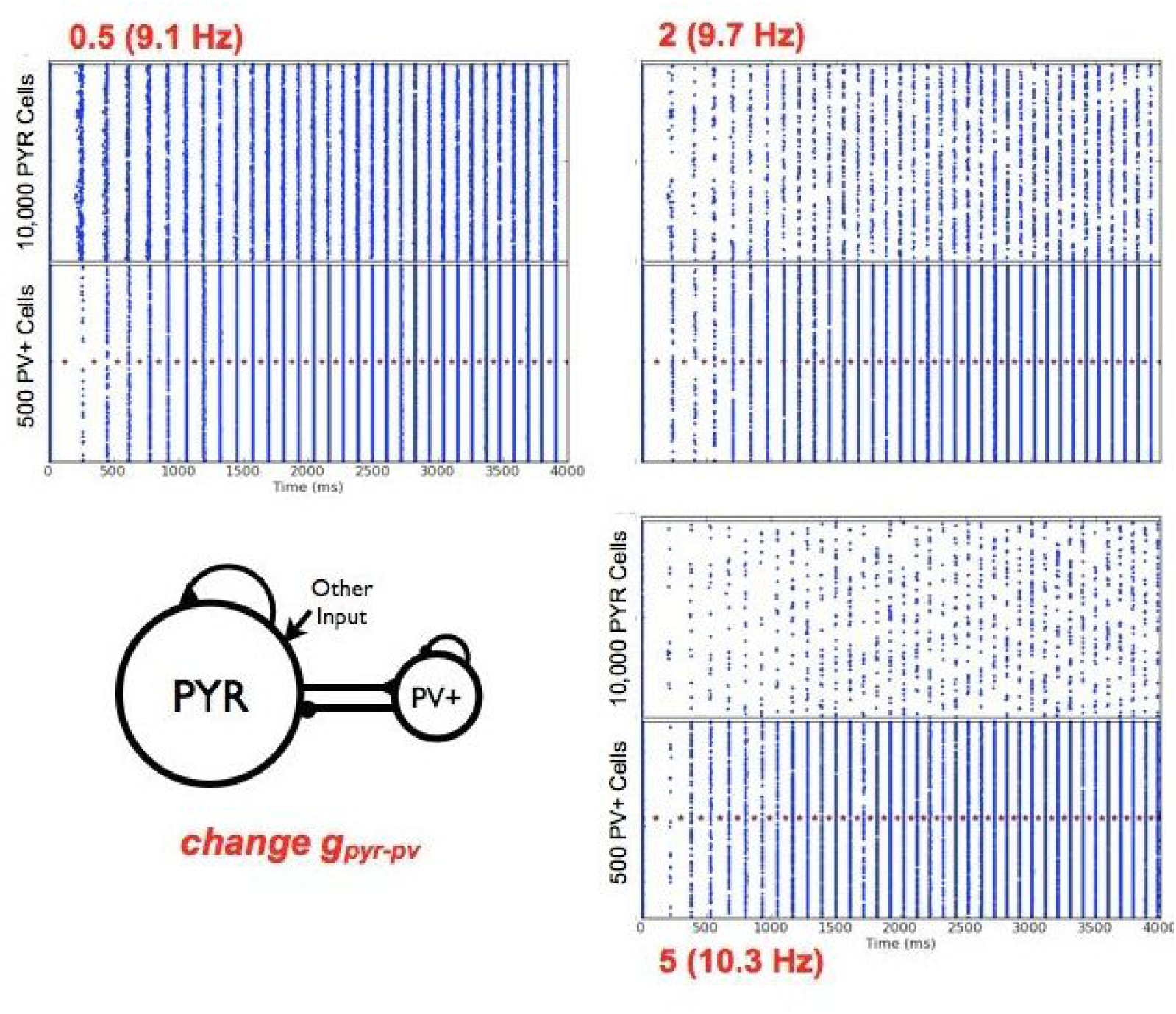
Changing Excitatory Conductance *g_pyr–pv_* from PYR to PV+ Cells. Three example raster plots are shown, with the *g_pyr–pv_* value shown in red with the network frequency as appropriate. The average number of cells per population burst is 290 (PYR) and 343 (PV+) for *g_pyr–pv_* = 0.5 nS; 81 (PYR) and 288 (PV+) for *g_pyr–pv_* = 2 nS. 32 (PYR) and 238 (PV+) for *g_pyr–pv_* = 5 nS. *Other parameter values: g_e,mean_, σ_e_, g_pyr_, g_pv–pyr_,c _PY R,PV,_c_PV,PYR_* = [0, 0.2, 0.084, 6,0.4, 0.5].

The particular parameter balances that allow the generation of theta population bursts affect not only the specific frequency of the population burst but also how robust it is, that is, how easily discernible it is (see Methods). This interdependence of cellular and synaptic properties affect how much the PYR and PV+ cell populations fire. It is clear that the addition of the PV+ cell population is what allows the PYR cell population to fire sparsely. However, how sparse the firing of PYR and PV+ cells are depends on where the parameter balance lies. From our simulations, we observe that as *c_PY R, PV_* decreases, the firing of PV+ cells becomes more sparse and PYR cells become less sparse within a theta population burst. This is illustrated by the box as described in the **Figure 7** bottom schematic. It makes sense that PV+ cells would fire less as they are receiving less excitatory drive with a reduced *c_PY R, PV_*. However, that PYR cells would fire less sparsely when the PV+ cells are firing less, indicates that how much the PYR cells fire is not *only* dependent on post-inhibitory rebound firing. There are clearly different balances going on. It is these different balances as brought forth from our thousands of simulations and analyses of them that allowed us realize that one could distinguish two scenarios by which theta rhythms emerge. Specifically, in Scenario A the PV+ cells fire less sparsely and the PYR cells fire more sparsely than in Scenario B. In **Figure 10**, we show an example of a Scenario A and a Scenario B theta frequency bursts. The different relative sparseness of PYR and PV+cells is apparent.

**Figure 10.**
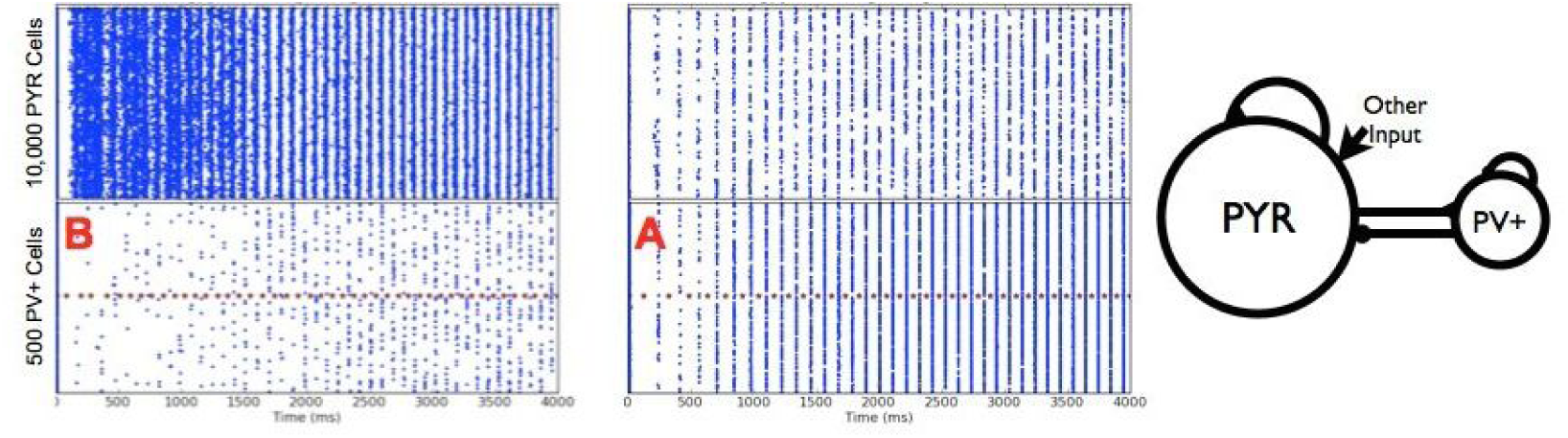
Theta Rhythm Generation by the two Different Scenarios. (*Left*) Scenario B: Same example as in *Figure 8* top. (*Right*) Scenario A: The average number of cells per population burst is 86 (PYR) and 185 (PV+). *Parameter values: g_pyr_,g_e,mean,_σ_e_, g_pyr–pv,_ g_pv–pyr_,c_Pv,PYR_,c_PY R,Pv_* = [0.084, 0, 0.2, 3, 8.7, 0.5, 0.2].

In **Table 4**, we present analyses regarding the average number of cells firing per burst as well as the average number of spikes per cell per burst for several parameter sets. From this table the two different scenarios can be appreciated from the relative firing of PYR and PV+ cells. Although it is clear that a PV+ cell spikes more than a PYR cell, that is, the relative comparisons are appropriate as observed in experiment, how active PYR and PV+ cells are is not completely in line with what exists in experiment, in terms of PV+ cells firing on each theta cycle. Scenarios A and B can be differentiated by how much of a role post-inhibitory rebound plays in the subsequent theta rhythm. In Scenario B, the PYR cells are less sparse and are less tightly bound to fire phase-locked to PV+ cells due to the post-inhibitory rebound, and the PV+ cell firing is more sparse. By contrast, in Scenario A, the PV+ cells fire more and post-inhibitory rebound plays a stronger role to more tightly control PYR cell phase-locking (so more tightly lined up with PV+ cells). These differences can be seen by comparing the cases shown in **Figure 10**.

**Table 4.** PYR-PV+ Cell Network Scenarios.

| Parameters<br>$g_{pyr}(nS), \sigma_e(nS)$<br>$g_{pyr-pv}(nS), g_{pv-pyr}(nS)$<br>$c_{PYR,PV}, c_{PV,PYR}$<br>$g_{e,mean} = 0nS$ | Burst<br>Freq<br>(Hz)<br>(from fft) | Average<br>Number<br>of Active<br>PYR cells<br>(/10,000)<br>per burst | Average<br>Number<br>of Active<br>PV+ cells<br>(/500)<br>per burst | Average<br>Number<br>of Spikes<br>(/PYR cell)<br>per 100<br>bursts | Average<br>Number<br>of Spikes<br>(/PV+ cell)<br>per 100<br>bursts |
| --- | --- | --- | --- | --- | --- |
| = [0.084, 0.2, 3, 8.7, 0.4, 0.5] | 10 | 53 | 273 | 1 | 55 |
| = [0.084, 0.2, 3, 8.7, 0.2, 0.5] | 9.7 | 86 | 185 | 1 | 37 |
| = [0.084, 0.2, 3, 8.7, 0.04, 0.5] | 8.9 | 300 | 79 | 3 | 16 |
| = [0.084, 0.2, 3, 8.7, 0.02, 0.5] | 8.3 | 522 | 54 | 5 | 11 |
| = [0.084, 0.2, 3, 8.7, 0.4, 0.3] | 10 | 53 | 249 | 1 | 50 |
| = [0.084, 0.2, 3, 8.7, 0.4, 0.7] | 10 | 54 | 280 | 1 | 56 |
| = [0.014, 0.6, 3, 8.7, 0.02, 0.3] | 11.7 | 548 | 22 | 2 | 8 |
| = [0.094, 0.6, 3, 8.7, 0.02, 0.3] | 12 | 514 | 34 | 5 | 7 |
| = [0.084, 0.2, 0.5, 6, 0.4, 0.5] | 9.1 | 290 | 343 | 3 | 69 |
| = [0.084, 0.2, 2, 6, 0.4, 0.5] | 9.7 | 81 | 288 | 1 | 58 |
| = [0.084, 0.2, 5, 6, 0.4, 0.5] | 10.3 | 32 | 238 | < .5 | 48 |

Excitatory and inhibitory currents for the chosen parameter sets in **Table 4** are shown in **Table 5**. Specifically, EPSCs and IPSCs to PYR and PV+ model cells are measured, and these values along with their ratios are given in **Table 5.** We note that there can be large EPSCs to PV+ cells and small EPSCs to PYR cells as observed experimentally. However, there is not always an appropriate match - IPSCs to PYR cells are too large and IPSCs to PV+ cells are too large in some cases. If current ratios rather than currents are compared, then it is always the case that EPSC/IPSC ratios are appropriate for PYR cells relative to experiment, but only in some cases are the EPSC/IPSC ratios for PV+ cells somewhat appropriate, that is, close to or greater than one. As such, we find that the EPSC/IPSC ratio to PV+ cells, but not the EPSC/IPSC ratio to PYR cells, allows a distinguishing between Scenarios A and B. We then conclude that Scenario B, but not Scenario A, is consistent with the experimental data, and so is the situation that is appropriate for the biological system. That is, one in which post-inhibitory rebound, although required to be present, plays less of a role in theta rhythm generation.

**Table 5.** PYR-PV+ Cell Network Scenarios - Currents.

| Parameters<br>$g_{pyr}(nS), \sigma_e(nS)$<br>$g_{pyr-pv}(nS), g_{pv-pyr}(nS)$<br>$c_{PYR,PV}, c_{PV,PYR}$<br>$g_{e,mean} = 0nS$ | EPSC<br>to<br>PYR<br>cell<br>(pA) | IPSC<br>to<br>PYR<br>cell<br>(pA) | E/I<br>Ratio<br>(PYR<br>cell) | EPSC<br>to<br>PV+<br>cell<br>(pA) | IPSC<br>to<br>PV+<br>cell<br>(pA) | E/I<br>Ratio<br>(PV+<br>cell) | Scenario |
| --- | --- | --- | --- | --- | --- | --- | --- |
| = [0.084, 0.2, 3, 8.7, 0.4, 0.5] | 4 | 1500 | << 1 | 700 | 180 | < 1 | A |
| = [0.084, 0.2, 3, 8.7, 0.2, 0.5] | 4 | 1600 | << 1 | 550 | 1300 | < 1 | A |
| = [0.084, 0.2, 3, 8.7, 0.04, 0.5] | 5 | 1500 | << 1 | 350 | 550 | $\approx 1$ | B |
| = [0.084, 0.2, 3, 8.7, 0.02, 0.5] | 7 | 730 | << 1 | 300 | 275 | $\approx 1$ | B |
| = [0.084, 0.2, 3, 8.7, 0.4, 0.3] | 4 | 2500 | << 1 | 650 | 1950 | < 1 | A |
| = [0.084, 0.2, 3, 8.7, 0.4, 0.7] | 4 | 1150 | << 1 | 740 | 1770 | < 1 | A |
| = [0.014, 0.6, 3, 8.7, 0.02, 0.3] | 1 | 410 | << 1 | 340 | 200 | > 1 | B |
| = [0.094, 0.6, 3, 8.7, 0.02, 0.3] | 7 | 430 | << 1 | 220 | 200 | $\approx 1$ | B |
| = [0.084, 0.2, 0.5, 6, 0.4, 0.5] | 7 | 2000 | << 1 | 480 | 2450 | < 1 | A |
| = [0.084, 0.2, 2, 6, 0.4, 0.5] | 4 | 2200 | << 1 | 650 | 2000 | < 1 | A |
| = [0.084, 0.2, 5, 6, 0.4, 0.5] | 4 | 2700 | << 1 | 870 | 1900 | < 1 | A |

## Discussion

### Summary, theta essense, explanation and predictions

We have developed microcircuit models and obtained an explanation for how theta rhythms can be generated in the hippocampus. We used a strategy, as schematized in **Figure 1**, in which we took advantage of a clear experimental context with developed mathematical models and theory for the context along with extensive parameter variation analyses. This computational analysis allowed us to differentiate between two scenarios of how theta rhythms could be generated, and only one of them is consistent with the experimental data. We suggest that spike frequency adaptation and post-inhibitory rebound in CA1 pyramidal cells are sufficient conditions (‘building blocks’) for the generation of theta rhythms with sparse excitatory cell firing. Moreover, if it is the case that spike frequency adaptation is required, then it is necessary to have post-inhibitory rebound. Further, our network simulations predict that theta rhythms are present when the input to the PYR cells has a non-zero fluctuating input conductance of 0.6 nS or less, and a zero mean conductance. Our network models are minimal but were able to capture an essence of the experimental data. As such, we consider our models as a foundation on which to build.

This building block understanding leads to the following claims and predictions. We first note that if PV+ cells are removed from the network, then there are no theta rhythms. This is known to be the case based on optogenetic silencing of PV+ cells by *Amilhon et al. (2015)*. We predict that spike frequency adaptation in PYR cells is required for theta rhythms to be present, as balanced by a large enough PYR cell network with connectivity that is minimal. The amount of spike frequency adaptation controls the existence of theta rhythms and its resulting frequency as built on our understanding from the theoretical mechanism. Thus, if cellular adaptation in PYR cells is selectively adjusted, say by modulating potassium and calcium-activated potassium channels, then one should see an effect on the frequency of theta rhythms, and with enough reduction in cellular adaptation, there would be no theta rhythms. Further, by selectively affecting either the amount of connectivity or the conductance from PYR to PV+ cells, the theta frequency would be strongly affected. However, if the amount of connectivity or the conductance from PV+ to PYR cells is selectively adjusted, then the theta frequency would be less affected. However, if this is reduced too much or removed, then there would be no theta rhythms as post-inhibitory rebound would not be present. Finally, from the large sets of parameter variations done and its correspondence with the data, we predict that theta rhythms in the hippocampus are generated via Scenario B (**Figure 7**). Although post-inhibitory rebound is still an important element to have theta rhythms (i.e., *c_PV, PYR_* cannot be zero) in Scenario B, it plays less of a role relative to Scenario A. If Scenario B is the mechanism underlying theta rhythms in the hippocampus, we predict that the probability of connections and/or conductance from PV+ to PYR cells is larger than the probability of connection and/or conductance from PYR to PV+ cells (see **Figure 7** schematic). It is of interest to note that large EPSCs onto PV+ cells can occur even if there is a minimally connected CA1 PYR cell network that sparsely fires. This is necessarily due to the large size of the PYR cell network.

Our explanation here of theta generation in the hippocampus has Aristotelian elements of efficient, material and formal causes (*Falcon, 2015)*. That is, we have identified building blocks (efficient cause) in our models of PYR and PV+ cell networks connected by inhibitory (*GABA_A_*) and excitatory (*AMPA*) synapses (material cause) in a mathematical model that requires parameter balances of cellular adaptation in a large PYR cell network with enough connectivity and postinhibitory rebound due to PV+ cells (formal cause). We note that a final cause is not considered in this work. A final cause would be considered as efforts to understand the function of theta rhythms. There are several ongoing efforts along these lines (see review in *Colgin (2013)*), but as emphasized by Colgin (2013) it is important to know what the underlying causes of theta generation are in the first place. In future work, for example, one might consider how our modeling work could be expanded and linked to functional theta modeling studies such as recent efforts by (*Chadwick et al., 2016*) to capture the flexibility of theta sequences by including phase precessing interneurons in septo-hippocampal circuitry.

### Related modeling studies

We have described previous mathematical modeling studies of theta rhythms (*Ferguson and Skinner, 2015*). Earlier models put forth theta generation mechanism ideas based on coherence between theta frequency firing in oriens-lacunosum/moleculare (O-LM) interneurons, in which a hyperpolarization-activated inward current (h-current) is critical for the coherence (*Rotstein et al., 2005; Wulff et al., 2009*). However, this has subsequently been shown to be unlikely since O-LM cells do not operate as theta pacemakers (*Kispersky et al., 2012*), an assumption in these earlier models.

In another hippocampal modeling study, it was shown that theta rhythms could be generated in a network of basket, O-LM and pyramidal cells (*Neymotin et al., 2011*). Although the focus of that study was not on theta generation per se, O-LM cells were strongly implicated in contributing to theta rhythms.

Based on recordings from the hippocampus of behaving rats, models were developed to support the observations that h-currents in pyramidal cells were needed to allow theta frequency spike resonance to occur (*Stark et al., 2013*). The mathematical models were focused on considering the contribution of h-currents in pyramidal cells, and did not specifically consider connectivity between pyramidal and parvalbumin-positive interneurons or the numbers of cells. This study emphasized the importance of post-inhibitory rebound in theta rhythms. Interestingly, this same study found that adaptation processes were not important contributing factors. In our modeling study, both cellular adaptation and post-inhibitory rebound are needed to bring about theta rhythms in the hippocampus. However, unlike Stark et al (2013)which had an experimental *in vivo* context, our models are based on a whole hippocampus preparation *in vitro,* thus removing contributions from other brain structures. In this way, we were able to focus on whether and how excitatory-inhibitory networks could produce theta rhythms considering network size, connectivity, and cellular characteristics. Given the complexity of theta rhythms, their modulation and induction by either fearful or social stimuli (*Tendler and Wagner, 2015*), one would expect there to be different balances of building blocks (cellular adaptation and post-inhibitory rebound) as well as additional ones underlying theta rhythms *in vivo.* However, it is considerably more challenging to dissect out an understanding of theta rhythm generation *in vivo.* Using our network models and mechanistic understandings derived from them, this challenge could be reduced.

A full-scale CA1 hippocampal model that includes many inhibitory cell types and gathered biological details (*Bezaire and Soltesz, 2013*) has recently been developed (*Bezaire et al., 2016*). It is loosely based on the whole hippocampus preparation and exhibits theta rhythms phase-locked with gamma oscillations as well as being able to show distinct phase relationships for different cell types. From model perturbations, it was determined that parvalbumin-expressing interneurons and neurogliaform cells and interneuronal diversity itself, are important factors in theta generation. Further, although not a particular focus, the generation of theta rhythms in these detailed network models required excitation levels that were neither too high or too low. Considering Aristotle’s four causes (*Falcon, 2015),* this descriptive understanding of theta generation can be considered as a material cause with elements of an efficient cause. However, due to its very detailed nature, it would be difficult to acquire a formal cause from it.

In a very recent study *Giovannini et al. (2017)* focused on the contribution of a non-specific cation (CAN) current in pyramidal cells as critical for the maintenance of theta oscillations in the isolated hippocampus preparation. Interestingly, similar to our work here, they showed that when pyramidal cells were coupled with inhibitory cells, theta oscillations became more robust. This work could be considered as a particular material cause explanation of theta rhythm generation.

### Spike frequency adaptation and post-inhibitory rebound

Spike frequency adaptation as a mechanism to generate populations bursts has been used before and in our previous work we sought to examine whether the amount of cellular adaptation expressed in pyramidal cells was appropriate to generate population bursts in pyramidal cell networks (*Dur-E-Ahmad et al., 2011; Ferguson et al., 2015a*). How best to model adaptation naturally depends on the questions being considered. For example, adaptation in signal transmission of sensory systems was examined as being due to either adaptation currents or dynamic thresholds (*Benda et al., 2010*). In another study, a fractional leaky integrate-and-fire model to capture spike frequency adaptation was developed to set a framework to help understand information integration in neocortex (*Teka et al., 2014*). For our CA1 pyramidal cell models, we used an Izhikevich type cellular model (*Izhikevich, 2006*) and the adaptation aspect was captured in the *d* and *a* parameters when fit to experimental frequency-current curves (*Ferguson et al., 2015b*). Further, for post-inhibitory rebound to be present in these models, the *b* parameter needed to be positive. Although the experimental data indicated that both strongly and weakly adapting pyramidal cells exist, this does not of course mean that there are simply two types, as the existence and amount of adaptation depends on the complement of biophysical ion channels in the cells, and what is uncovered by the particular experimental protocol. In the work here, we used both types of pyramidal cell models, but networks of weakly adapting cells only were not able to produce theta rhythms given other constraints of the experimental context.

Post-inhibitory rebound inhibition exists in CA1 pyramidal cells as recorded in this experimental preparation (*Ferguson et al., 2015b; Goutagny et al., 2009*). H-currents in pyramidal cells clearly play a key role in their ability to express post-inhibitory rebound. However, not only the presence, but also the distribution of these currents together with the distribution of other currents need to be taken into consideration. Specifically, it was shown that post-inhibitory rebound is rarely observed in physiological conditions unless unmasked by the blocking of A-type potassium currents (*Ascoli et al., 2010*). Both h-currents and A-type potassium currents are known to have a non-uniform distribution along pyramidal cell dendritic arbors, and putative functional contributions of this to temporal synchrony have been made (*Vaidya and Johnston, 2013*). It is interesting to note that a difference in the dorsal to ventral patterning of h-currents has been found (*Dougherty et al., 2013*), bringing to light another possibility of how theta rhythms could be modulated. Further, travelling theta waves have been observed in both rodent (*Lubenov and Siapas, 2009; Patel et al., 2012*) and human hippocampus (*Zhang and Jacobs, 2015*) suggesting a coupled oscillator organizational motif in the hippocampus. Although a *‘weakly* coupled oscillator’ terminology has been invoked in describing these waves (*Colgin, 2013*), this should not be confused with the mathematical theory where the assumption of weakly coupled oscillators is used to reduce the system to a phase-coupled system that is easier to analyze (*Schwemmer and Lewis, 2012*).

### Limitations

Given our highly simplified and minimal network models we did not expect to find a perfect matching to the experimental data. However, it is important to note that given our minimal models, we were able to examine several thousand parameter sets which in turn enabled us to explore and understand what balances might be important in bringing about theta rhythms. This ‘balance’ and ‘building block’ understanding can serve as a basis for how theta rhythm frequency and existence can be modulated by additional inputs from other brain regions as well as modulation that would affect adaptation and post-inhibitory rebound.

Our network models are minimal, but they were able to produce theta rhythms with sparse firing, as represented by population bursts, allowing us to suggest sufficient and necessary conditions for their generation. While our models took into consideration network size, connectivity, and cellular characteristics in a clear experimental context, architecture was not considered. That is, connectivity used in the network models was random. This is clearly overly simple relative to experiment, especially considering the recent finding of motifs in pyramidal cells in the CA3 region of hippocampus (*Guzman et al., 2016*). However, it is a reasonable first approximation which allowed us to explore a wide expanse of connectivities.

Variability in intrinsic cell properties and a mixing of weakly and strongly adapting pyramidal cells could be considered. Introducing variability immediately raises the question of how it should be done. For example, work in *Harrison et al. (2015)* considered the cellular heterogeneity of cortical cells, but the goal of that study was focused on determining how to capture a wide range of experimental heterogeneity of cortical cells in simple models, and was not focused on particular network dynamics or rhythms per se. However, given our large explorations of conductances, connectivities and noisy input, variability to a certain extent was present in our simulations.

Further, only one type of inhibitory cell was included in our networks, that is, the fast-firing parvalbumin-positive cell type. It is unlikely that only this one type of inhibitory cell contributes to theta rhythms, but we identified this as a good place to start given that *Amilhon et al. (2015*) found that they were an essential requirement. By no means do our models imply that other inhibitory cell types are unimportant. On the contrary, since our models clearly do not fully capture the experimental data (e.g., model PV+ cells fire too sparsely relative to experiment), aspects are clearly missing. Given this limitation, it would be surprising if our models were able to mimic all aspects of the experiment. Since PV+ fast-firing cells also include bistratifed and axo-axonic cells, an expansion of PV+ cell networks along these lines could be considered, building on our earlier work when theta rhythms were imposed based on experimental EPSCs (*Ferguson et al., 2015c*). Given the diversity of inhibitory cell types (e.g., see *Chamberland and Topolnik (2012)* and the different types of PV+ cells (*Baude et al., 2007*), we did not specifically try to scale the number of PV+ cells as we did for the pyramidal cells. However, since we fully explored connectivity ranges between PV+ and pyramidal cells, this was in effect included. The inclusion of O-LM cells and other inhibitory cell types in the network models is important moving forward to be able to understand how they modulate theta rhythms (*Amilhon et al., 2015; Leao et al., 2012; Sekulic and Skinner, 2017*).

### Conclusions and future work

In summary, although limited, we think that the present network models represent a reasonable skeleton since they are able to capture the experimental data in various ways. At the end of the day, it is always a balance. Our network models represent closed, self-consistent, accessible models that can generate theta rhythms in hippocampus CA1. We intend it as a starting point on which to build in understanding theta rhythms in the hippocampus. Specifically, the network model building block balances need to be fully analyzed so that a more solid ‘formal cause’ of explanation can be obtained, beyond what was obtained from the computational, parameter variation analyses done here. Further, in combination with full-scale models such as *Bezaire et al. (2016)* it may be possible to obtain an understanding that can fully encompass efficient, material and formal causes in the Aristotelian sense, and which could subsequently help in a final cause understanding.

Overall, our models can serve as a backbone on which other cell types as well as details of particular cell types (biophysical channels, dendrites and spatial considerations), modulatory effects, input from medial septum can be incorporated. However, in doing this, it is important to note that interaction and testing with experiment should be designed accordingly, given the strategy used in developing our models (**Figure 1**).

## Acknowledgments

We thank NSERC and CIHR of Canada for funding. Many thanks to Sylvain Williams and his lab for open and friendly collaborations over the years - we have learnt a lot! Special thanks to Vladislav Sekulić and Sue Ann Campbell for their detailed suggestions, and to all lab members for their comments and feedback.

